# dsMTL - a computational framework for privacy-preserving, distributed multi-task machine learning

**DOI:** 10.1101/2021.08.26.457778

**Authors:** Han Cao, Youcheng Zhang, Jan Baumbach, Paul R Burton, Dominic Dwyer, Nikolaos Koutsouleris, Julian Matschinske, Yannick Marcon, Sivanesan Rajan, Thilo Rieg, Patricia Ryser-Welch, Julian Späth, The COMMITMENT consortium, Carl Herrmann, Emanuel Schwarz

## Abstract

Multitask learning allows the simultaneous learning of multiple ‘communicating’ algorithms. It is increasingly adopted for biomedical applications, such as the modeling of disease progression. As data protection regulations limit data sharing for such analyses, an implementation of multitask learning on geographically distributed data sources would be highly desirable. Here, we describe the development of dsMTL, a computational framework for privacy-preserving, distributed multi-task machine learning that includes three supervised and one unsupervised algorithms. dsMTL is implemented as a library for the R programming language and builds on the DataSHIELD platform that supports the federated analysis of sensitive individual-level data. We provide a comparative evaluation of dsMTL for the identification of biological signatures in distributed datasets using two case studies, and evaluate the computational performance of the supervised and unsupervised algorithms. dsMTL provides an easy-to-use framework for privacy-preserving, federated analysis of geographically distributed datasets, and has several application areas, including comorbidity modeling and translational research focused on the simultaneous prediction of different outcomes across datasets. dsMTL is available at https://github.com/transbioZI/dsMTLBase (server-side package) and https://github.com/transbioZI/dsMTLClient (client-side package).

## Introduction

The biology of many human illnesses is encoded in a vast number of genetic, epigenetic, molecular, and cellular parameters. The ability of Machine Learning (ML) to jointly analyze such parameters and derive algorithms with potential clinical utility has fueled a massive interest in biomedical ML applications. One of the fundamental requirements for such ML algorithms to perform well is the availability of data at a large scale, a challenge of steadily declining importance due to the ever-increasing availability of biological data^1–3^. As data can often not be freely exchanged across institutions due tothe need for protection of the individual privacy, the utility of ‘bringing the algorithm to the data’ is becoming apparent. Technological solutions for this task have thus risen in popularity and exist in various forms. One of the most straightforward approaches is the so-called federated ML, where algorithms are simultaneously learned at different institutions and optimized through a privacy-preserving exchange of parameters. Other approaches for this task include the training of ML algorithms on temporarily combined data stored in working memory^4^ or the more recently introduced ‘swarm-learning’ approach^5^. One commonality of most ML algorithms, federated or not, is the assumption that all investigated observations (e.g. illness-affected individuals) represent the same underlying population. However, in biomedicine, this is rarely the case, as biological and technological factors frequently induce cohort-specific effects that limit the ability to identify reproducible biological findings. Multitask Learning (MTL) can address this issue through the simultaneous learning of outcome (e.g. diagnosis) associated patterns across datasets with dataset-specific, as well as shared, effects. Multi-task learning has numerous exciting application areas, such as comorbidity modeling, and has already been applied successfully for e.g. disease progression analysis^6^.

Here, we describe the development of dsMTL (‘Federated Multi-Task Learning for DataSHIELD’), a package of the statistical software R, for **Fe**derated **M**ulti-**T**ask **L**earning (FeMTL) analysis (**Figure 1**). dsMTL was developed for DataSHIELD^7^, a platform supporting the federated analysis of sensitive individual-level data that remains stored behind the data owner’s firewall throughout analysis^8^. dsMTL includes three supervised and one unsupervised federated multi-task learning algorithms that extend algorithms previously developed for non-federated analysis (for R implementations, see ^9,10^). Specifically, the **dsMTL_L21** approach allows for cross-task regularization, building on the popular LASSO method, in order to identify outcome-associated signatures with a reduced number of features shared across tasks. The non-federated version of this approach has previously been applied to simultaneously predict multiple oncological outcomes using gene expression data^11^. The **dsMTL_trace** approach constrains the coefficient vectors in a low-dimensional space during the training procedure to penalize the complexity of task relationships, resulting in an improved generalizability of the models. In a non-federated implementation, this method has previously been used to predict the response to different drugs, and the identified models showed a high degree of interpretability in the context of the represented drug mechanism^12^. **dsMTL_net** incorporates the task relationships that can be described as a graph, in order to improve biological interpretability. In a non-federated version, this technique has previously been used for the integrative analysis of heterogeneous cohorts^13^ and for the prediction of disease progression^14^. The **dsMTL_iNMF** approach is an unsupervised, integrative non-negative matrix factorization method that aims at factorizing the cohorts’ data matrices into shared and dataset-specific components. Such modeling has been applied to explore dependencies in multi-omics data for biomarker identification^10,15^. In addition to the FeMTL methods, we also implemented a federated version of conventional Lasso (dsLasso) ^16^ in dsMTL package due to its wide usage in biomedicine and as a benchmark for testing the performance of the federated MTL algorithms.

**Figure 1.**
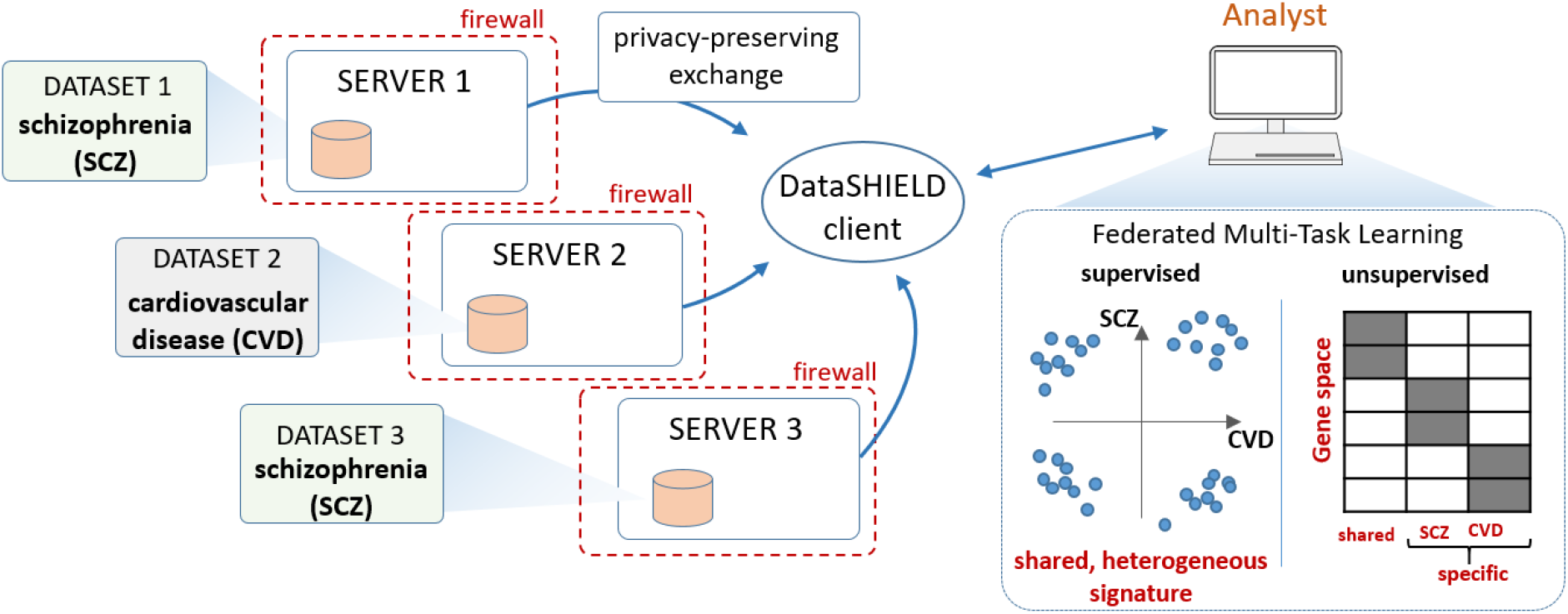
Schematic illustration of dsMTL using comorbidity modeling of schizophrenia and cardiovascular disease as an example. Multiple datasets stored at different institutions are used as a basis for federated MTL. dsMTL was developed in the DataSHIELD ecosystem, which provides functionality regarding data management, transmission and security. Data are analyzed behind a given institution’s firewall and only algorithm parameters that do not disclose personally identifiable information are exchanged across the network. dsMTL contains algorithms for supervised and unsupervised multi-task machine learning. The former aims at identifying shared, but potentially heterogeneous signatures across tasks (here, diagnostic classification for schizophrenia and cardiovascular disease). Unsupervised learning separates the original data into shared and cohort-specific components, and aims at revealing the corresponding outcome-associated biological profiles.

To explore the utility of the dsMTL algorithms, we used a network comprising three servers. These servers hosted simulated data with variable degrees of cross-dataset heterogeneity, in order to test the ability of the MTL algorithms to suitably characterize shared and specific biological signatures. In addition, we analyzed actual RNA sequencing and microarray data across the three-server network, to show that the accurate analysis can be performed in acceptable runtime using dsMTL in real network latency.

## Results

Here we show the results for two case studies. The first case study aims at demonstrating the utility of the supervised **dsMTL_L21** algorithm to identify ‘heterogeneous’ target signatures across the data network. With ‘heterogeneous’ we describe signatures that involve the same features (e.g. genes) but with potentially differing signs (indicating differential directions of influences) across datasets. In contrast, ‘homogeneous’ signatures relate to the same features and signs across datasets. The second case study focuses on the unsupervised **dsMTL_iNMF** method and explores the utility of the federated implementation, compared to the aggregation of local NMF models, to disentangle shared and cohort-specific components across datasets. For all case studies, we evaluated the signature identification accuracy as the major metric. For predictions of clinical outcomes, the prediction accuracy was also demonstrated.

### Case study 1 – distributed MTL for identification of heterogeneous target signatures

With the aim to identify ‘heterogeneous’ signatures, we compared the performance of dsMTL_L21, dsLasso and the bagging of glmnet models. As part of this, we explored the sensitivity of these methods to different sample sizes (*n*) relative to the gene number (*p*). **Figure 2** shows the resulting prediction performance and gene selection accuracy, each averaged over 100 repetitions. dsLasso showed the worst prediction performance in this heterogeneous setting, and dsMTL_L21 slightly outperformed the aggregation of local models (glmnet). Similarly, the gene selection accuracy of dsLasso was inferior to that of dsMTL_L21 and glmnet-bagging, which showed similar performance when the sample size is sufficiently large, e.g. the number of subjects approximately equal to the number of genes (n/p ^~^1). However, with a decreasing n/p ratio, dsMTL_L21 showed an increasing superiority over the other methods, especially for n/p=0.15, where the gene selection accuracy of dsMTL_L21 was over 2.8 times higher than that of the bagging technique.

**Figure 2.**
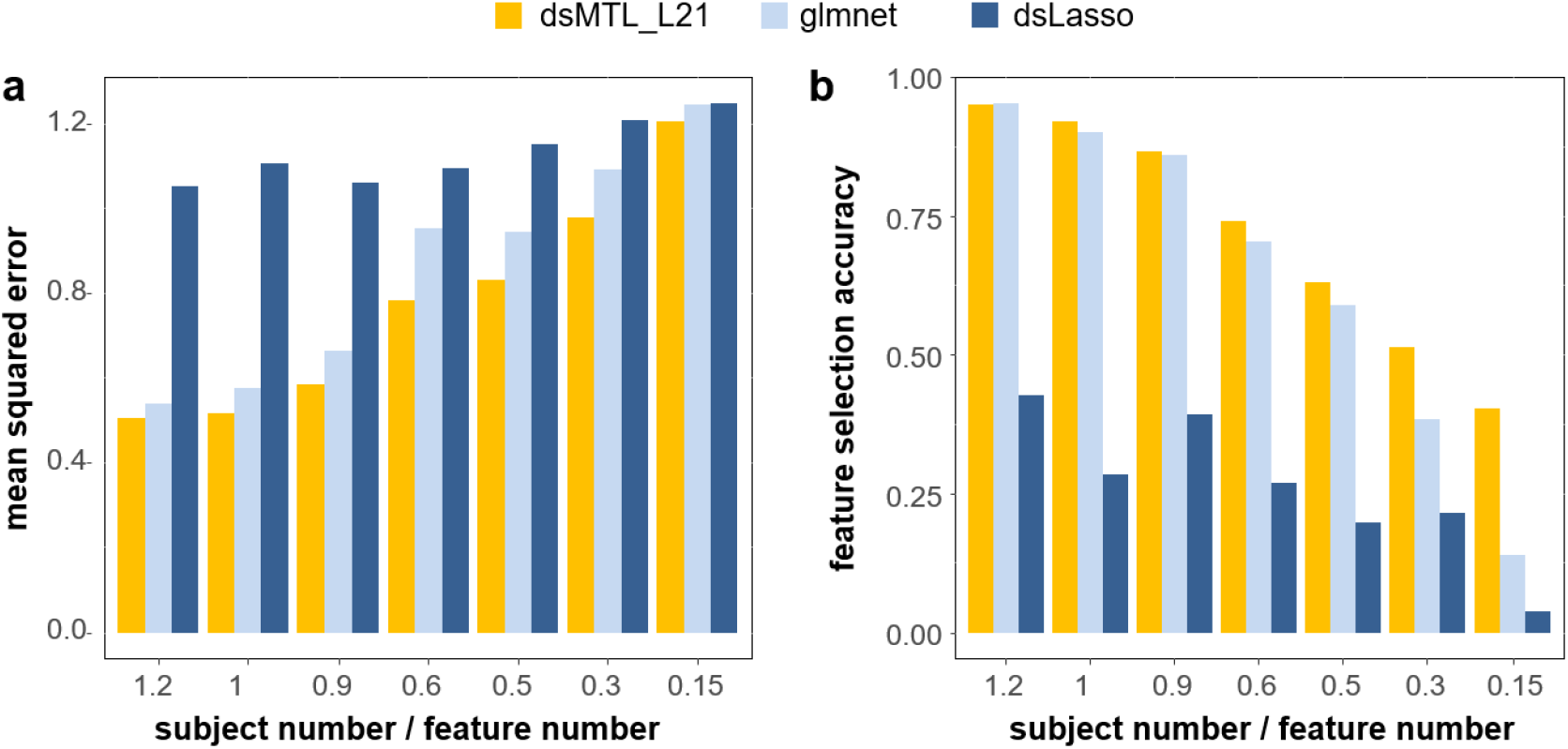
Analysis of ‘heterogeneous’ signatures of continuous outcomes in simulated data stored on three servers. The figure shows the **a)** prediction accuracy expressed as the mean squared error and **b)** the feature selection accuracy for different subject/feature number ratios. The respective values were averaged across the three servers, and across 100 repetitions, in order to account for the effect of sampling variability.

### Case study 2 – distributed iNMF for disentangling shared and cohort-specific signatures

**Figure 3** shows the performance of distributed and aggregated local NMF methods for disentangling shared and cohort-specific signatures from multi-cohort data, given different ‘severities’ of the signature heterogeneity. For both types of signatures, dsMTL_iNMF outperformed the ensemble of local NMF models for any heterogeneity severity setting. Notably, even with increasing heterogeneity, the accuracy of dsMTL_iNMF to capture shared genes remained stable at approximately 100%, illustrating the robustness of dsMTL_iNMF against the heterogeneity’s severity shown in **Figure 3c**. In contrast, for the ensemble of local NMF, the gene selection accuracy of the shared signature continuously decreased to approximately 50% (20% of outcome-associated genes were shared among cohorts), while the gene selection accuracy of cohort-specific signatures continuously increased to 75% (20% of outcome-associated genes were shared among cohorts) as shown in **Figures 3a** and **3b**.

**Figure 3.**
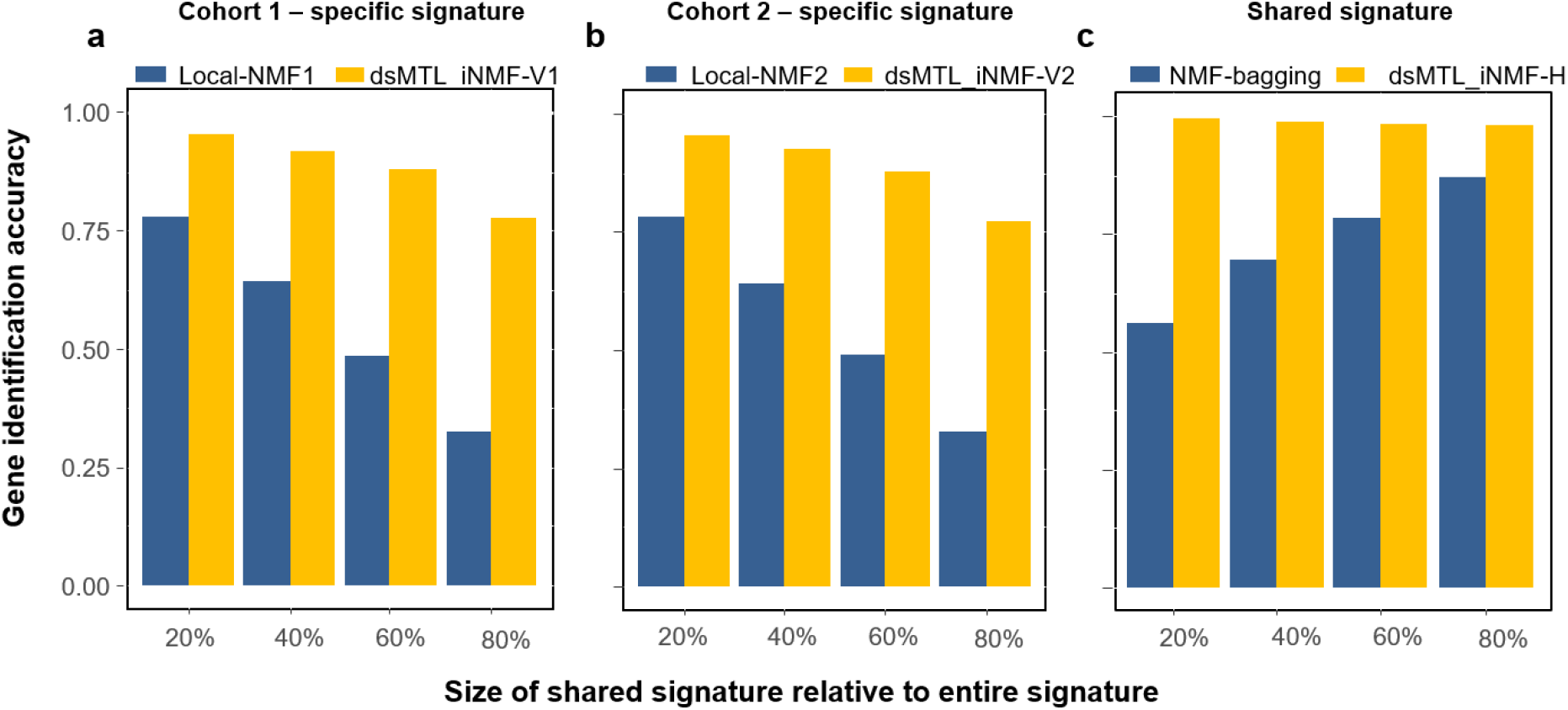
The gene identification accuracy for shared and specific signatures using simulated data. **a)** the identification accuracy of important genes for cohort 1. **b)** the identification accuracy of important genes for cohort 2. **c)** the identification accuracy of genes comprised in the shared signature. Local-NMF1 and Local-NMF2 were the cohort-specific gene sets identified by local NMF, which were combined into “NMF-bagging” for the shared gene set. dsMTL_iNMF-H was the predicted shared gene set using dsMTL_iNMF. dsMTL_iNMF-V1 and dsMTL_iNMF-V2 were the predicted cohort-specific gene sets identified using dsMTL_iNMF (see Supplementary Figure 1). The proportion of genes harbored by the shared signature was varied from 20% to 80% illustrating the impact of the heterogeneity severity. The model was trained using rank=4 as model parameter. The results for a broader spectrum of rank choices can be found in **Supplementary Figure 2** illustrating that the superior performance of dsMTL_iNMF was not due to the choice of ranks.

### Efficiencyof supervised dsMTL

We aimed at determining the efficiency of supervised dsMTL using the real molecular data and the actual latency of a distributed network. Using a three-server scenario (see **Table 2 Supplementary Results**; two servers at the Central Institute of Mental Health, Mannheim; one server at BioQuant, Heidelberg University) we analyzed four case-control gene expression datasets of patients with schizophrenia and controls (median n=80; 8013 genes). **Supplementary Table 3** shows the comparison between dsLasso and mean-regularized dsMTL_net, which were trained (cross-validation + training) and tested in approximately 8min and 10min, respectively, with the time-difference being due to the increased network access of dsMTL. The prediction accuracy of dsMTL was slightly higher than that of dsLasso, consistent with our previous study^13^. Regarding model interpretability, dsLasso captured a signature comprising 38 genes but could not distinguish shared and cohort-specific effects. Mean regularized dsMTL identified a signature with 10 genes shared among all cohorts, with 163 genes shared by two cohorts, as well as three cohort-specific signatures comprising 1532 genes.

### Efficiency of unsupervised dsMTL

The cohorts and server information is shown in **Supplementary Table 4**. It took 34.9 minutes (1,003 times network accesses) to train a dsMTL_iNMF model with 5 random initializations (^~^7 min for each initialization). The factorization rank k=4 was selected as the optimal parameter. In **Supplementary Figure 1**, the objective curve illustrates that the training time was sufficient for model convergence. In this analysis, a shared signature comprising 473 genes between SCZ and BIP was identified, while two disease-specific signatures containing 37 genes for SCZ and 152 genes for BIP, respectively, were found.

## Discussion

We here present dsMTL – a secure, federated multi-task learning package for the programming language R, building on DataSHIELD as an ecosystem for privacy-preserving and distributed analysis. Multi-task learning allows the investigation of research questions that are difficult to address using conventionalML, such as the identification of heterogeneous, albeit related, signatures across datasets. The implementation of a privacy-preserving framework for the distributed application of MTL is an essential requirement for the large-scale adoption of MTL. Using such a distributed server setup, we demonstrate the applicability and utility of dsMTL to identify biomarker signatures in different settings. For applications where the target biomarker signatures are different, but relate to an overlapping set of features (explored here as the ‘heterogeneous’ case), conventional machine learning would not be a meaningful algorithm choice. We show that MTL is able to identify the target signatures with high confidence and may thus be a reasonable choice for a diverse set of interesting analyses. As mentioned above, a particularly noteworthy application is comorbidity modeling, where the target signatures index the shared (although potentially heterogeneously manifested) biology of multiple, clinically comorbid conditions. Such analyses could potentially be a powerful, machine learning-based extension of comorbidity modeling approaches based on univariate statistics that have already been very useful for characterizing the shared biology of comorbid illness^17^. We show that unsupervised MTL can disentangle the shared from cohort-specific effects, demonstrating its potential utility for comorbidity analysis. Other applications for this method include the analysis of biological patterns shared across clinical symptom domains, between clinical and demographic characteristics, or with digital measures, such as ecological momentary assessments.

The use of dsMTL follows the concept of the so-called “freely composing script” in the DataSHIELD ecosystem. It organizes a given dsMTL workflow as a free composition of dsMTL, DataSHIELD, and local R commands (e.g. R base functions, customer-defined functions and CRAN packages) into a script, such that the geo-distribution of datasets and the federated computation are transparent to users. This concept is similar to that of the “freely composing apps” used in a recently presented federated ML application^18^, which allows flexible scheduling of functions in the form of apps and improves the federated data analysis flexibility for users. In addition to dsMTL, other packages in the DataSHIELD ecosystem exist for e.g. “big data” storage and management^19^, various statistical tests^7,19^ and deep learning^19,20^.

Interesting future developments of the dsMTL approach could include the implementation of asynchronous communication, which provides a probabilistically approximate solution but faster convergence^21,22^. Furthermore, integration of other popular systems for ML, such as tensorflow^23^, for which interfaces with the R language already exist, would provide valuable additions to the DataSHIELD system. Finally, a noteworthy consideration is an architecture underlying the distributed data infrastructure. DataSHIELD builds on a centralized (“client-server”) architecture and each data provider needs to install a well-configured data warehouse. Such infrastructure is suitable for long-term collaboration scenarios and large consortia projects that conduct a broad spectrum of complex analyses requiring high flexibility. However, in other scenarios that require more temporary and easy-compute collaboration setups, a server-free or decentralized architecture^24^ might be more suitable, because the cost of data provider for participating is low.

In conclusion, the dsMTL library for the programming language R provides an easy-to-use framework for privacy-preserving, federated analysis of geographically distributed datasets. Due to its ability to disentangle shared and cohort-specific effects across these datasets, dsMTL has numerous interesting application areas, including comorbidity modeling and translational research focused on the simultaneous prediction of different outcomes across datasets.

## Methods

### Modeling

All methods part of dsMTL share the identical form,

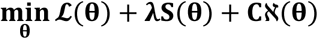

where 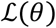 is the data fitting term (or loss function), the major determinant of the solutions obtained from model training. 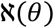 and *S*(*θ*) are the penalties of *θ* with the aim to incorporate the prior information. 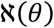 is a non-smooth function and able to create sparsity, while *S*(*θ*) is smooth. *λ* and *C* are the hyper-parameters to control the strength of the penalties. More technical details can be found in the supplementary methods.

In dsMTL, two approaches for sharing information across cohorts are included, 1) shared parameters and 2) cross-task regularization, leading to a slightly different distributed computation. The shared parameters are estimated using all cohorts. For cross-task regularization, the cohort-specific parameters are estimated using only the local data, and then tuned by considering parameters from other cohorts.

### Efficiency

Most dsMTL methods aim at training an entire regularization tree. The determination of the λ sequence controls the tree’s growth and is essential for computational speed. The λ sequence should be accurately scaled to both capture the highest posterior and avoid overwhelming computations. Inspired by a previous study^25^, we estimate the largest and smallest λ from the data by characterizing the optima of the objective using the first-order optimal condition and then interpolate the entire λ sequence on a log scale (see supplementary methods for more details). In addition, several options are provided to improve the speed of the algorithms by decreasing the precision of the results, i.e., 1) the number of digits of parameters for transformation can be specified to reduce the network latency; 2) several termination rules are provided, some of which are relaxed; 3) the depth of the regularization tree can be shortened. More details can be found in supplementary methods.

Besides the efficiency of the federated ML/MTL methodology, the import/export of “big data” cohorts is also crucial for computational efficiency, where e.g. uncompressed GWAS data requires tens of gigabytes, leading to time-consuming data import. dsMTL was designed to support a wide variety of data types. For this, an architecture package resourcer^19^ developed by the DataSHIELD community was incorporated to facilitate the efficient import and export of large-scale datasets in compressed formats. For example, in DataSHIELD, GWAS data of the PLINK file formats can be read and processed using the software PLINK^26^ as the backend^19^.

### Security

dsMTL was developed based on DataSHIELD^8^, which provides comprehensive security mechanisms not specific to machine learning applications. For example, 1) DataSHIELD requires the data analysis to only occur behind the firewall; 2) each server is only allowed to communicate with a set of clients with fixed IP addresses; 3) the network communication is protected by an SSL protocol; 4) an R parser^8^ implemented on the server rejects the calling of unwanted functions; and 5) the so-called ‘disclosure control’^8^ on the server ensures that the returned response does not contain any disclosive information.

In addition, several permissions can be set by the data providers to fully control the usage of their data. These permissions describe the degree of accessibility of data and functions on the server i.e. “*which users* can perform *what actions* on *what data*”. In an extremely secure example, a user could be granted to check the summary of a given dataset but cannot perform any actions because no functions were granted. With these settings, DataSHIELD allows customizing the security protection strategies according to the specific requirements of the applications. For statistical and machine learning analyses, DataSHIELD assumes that summary statistics are safe to share.

dsMTL inherits all these security mechanisms. In addition, we considered potential ML-specific privacy leaks, such as membership inference attacks^27^ and model inverse attacks^28^. Inverse attacks aim at extracting the individual observation-level information from the models. Membership inference attempts to decide if an individual was included in a given training set using the model. All these techniques require a complete model for inference. Since multi-task learning returns multiple matrices, returning an incomplete model could be one strategy against these attacks. For example, dsMTL_iNMF in dsMTL only returns the homogenous matrix (H), whereas the cohort-specific components (*V_k_, W_k_*) never leave the server. For example, in a two-server scenario, one (H) out of five output matrices is transmitted between the client and the servers. With such an incomplete model, inverse construction of the raw data matrix becomes difficult, and the risk of an inverse attack and membership inference is reduced. For most biomedical analyses, the H matrix is sufficient for subsequent studies. In addition, if the analyst was authorized to access the raw data of the server, the so-called “data key mechanism” (see supplement) would allow the analyst to retrieve all component matrices. For supervised multi-task learning methods in dsMTL, all models have to be aggregated within the clients, and thus we suggest the data providers enable the option on the server that rejects a returned coefficient vector containing parameter numbers exceeding the number of subjects. In this way, the model is not saturated and more robust to an inverse attack.

### Proof of concept with simulation and actual data

Two case studies and speed-tests were conducted to demonstrate the suitability of dsMTL methods to analyze heterogeneous cohorts, compared to federated ML methods and ensemble of local models regarding the prediction performance, interpretability and computational speed. An overview of methodological aspects related to the case studies is detailed below. For an extensive methodological description, please see the supplementary Methods.

#### Case study 1

In this case study, the heterogeneous cohorts were generated with the same set of outcome-associated genes. These however showed different directionality of their respective associations with the outcome. A three-server scenario was simulated. 150 out of 500 features with random signs across cohorts were simulated. Seven tests were created for simulating different n/p 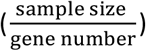 ratios. The n/p ratio was {1.2,1,0.9,0.6,0.5,0.3,0.15} with the number of subjects {600,500,450,300,250,150,75} for each test. 500 genes were created for each server. The test sample consisted of 200 subjects for each server. Data were generated as follows:

Given gene number p = 500, the models of three cohorts were {*w*^(1)^, *w*^(3)^, *w*^(3)^} where *w*^(.)^ = p × 1. A shared signature comprising 150 genes was generated for each *w*^(.)^ but with random signs, 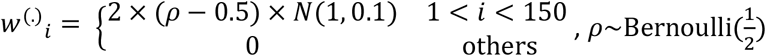. The expression values of each subject across cohorts were generated as x = 1 × p where *x_j_*~*N*(0,1). The numeric outcome (e.g. symptom severity) y = xw^(*i*)^ in cohort i was standardized in a normal distribution *N*(0,1), then model-irrelevant noise with 50% of the variance of the true signal was added y = y + *N*(0, 0.5).

dsMTL_L21 and dsLasso were trained as the federated learning system, and the hyper-parameter was selected using 10 fold in-cohort cross-validation. For glmnet, the ensemble technique was only applied on the gene selection due to the consistent gene set of their signatures. The mean squared error (mse) was used as the measure of prediction performance. To account for the sampling variance, we repeated each analysis 100 times.

#### Case study 2

In this case study, two heterogeneous RNA-seq cohorts were created to simulate a comorbidity analysis, where the genes were separated to be part of either a shared signature among cohorts, cohort-specific signatures or diagnosis-unassociated genes. The dsMTL_iNMF was compared to the ensemble of local NMF regarding the selection accuracy of shared/cohot-specific genes, in particular impacted by the severity of heterogeneity. Here the severity of heterogeneity refers to the proportion of the genes harbored by the shared signature over all diagnosis-associated genes. The data simulation protocol for RNA-seq data can be found in the **Supplementary Methods**.

A two-server scenario was simulated. As shown in **Supplementary Table 1**, for the data of each server, 1000 genes and 200 subjects were simulated, 50% of the genes were diagnosis-unassociated and the remaining genes were part of the disease signature. The genes comprised by shared signatures were identical for data of two servers, and the genes comprised by cohort-specific signatures did not overlap. The case-control ratio was balanced for each server. Four tests were performed by varying the proportion of genes in the shared signature over all diagnosis-associated genes from 20% to 80%.

The training of dsMTL_iNMF results in three outputs related to the original input data: the shared gene ‘exposure’ (H), cohort-specific gene ‘exposure’ (V) and sample ‘exposure’ (W). We measured the association between the sample exposure and the diagnosis as the weight of each latent factor. The shared(or specific) gene signature was identified as the weighted summation of the shared (or specific) gene exposures over latent factors. To quantify the important genes related to a given signature, we binarized the gene signature according to the mean (0-1 vector, values larger than the mean were assigned). To assess the performance of the gene identification, we associated the selected genes set with the ground truth (0-1 vector, signature genes were 1). The assessment was applied to shared and cohort-specific genes in parallel. Based on this metric, three gene sets were derived as output from dsMTL_iNMF, called dsMTL_iNMF-H, dsMTL_iNMF-V1 and dsMTL_iNMF-V2, and these related to the shared, cohort 1 specific and cohort 2 specific gene signature, respectively. The same strategy was applied to analyze the ensemble of local NMF models. For each cohort, the specific gene signature was the weighted summation of gene exposure over latent factors, and then binarized as the specific gene set (called local-NMF1 and local-NMF2). The shared gene signature was identified as the sum of the specific gene signature over cohorts, and then binarized as the shared gene set(NMF-bagging). We then compared 1) NMF-bagging and dsMTL_iNMF-H for the accuracy related to the isolation of shared genes; 2) dsMTL_iNMF-V1 and local-NMF1 as well as dsMTL_iNMF-V2 and local-NMF2 for the accuracy of isolating cohort-specific genes.

#### Computational speed of supervised dsMTL

We aimed at identifying the efficiency of supervised dsMTL using real molecular data and given the real network latency. Four independent schizophrenia case-control cohorts were used for this analysis. The training cohorts consisted of three datasets comprising prefrontal cortex gene expression data (available from the GEO repository under accession numbers GSE53987, GSE21138 and GSE35977). A detailed description of these datasets can be found in their respective original publications^29–31^. The dataset used for algorithm testing was from the HBCC (n=422) cohort comprising genome-wide gene expression data quantified by microarray (dbGAP ID: phs000979.v3.p2). A detailed description of this dataset can be found in the original publication^32^. As shown in **Supplementary Table 2**, three servers were used for training algorithms. Two servers were held at the Central Institute of Mental Health, Mannheim while the third was positioned at the BioQuant institute, Heidelberg.

Using this data, we repeated a previously described analysis^13^, in order to evaluate computational speed in a federated analysis setting. Here we show the formulation of the mean regularized MTL using dsMTL_net:

The cohort-level batch effect was assumed to be Gaussian noise affecting the true coefficient of gene i and cohort j *w_ij_* = *w_i_* + *ϵ_j_*, *ϵ_j_* ∈ *N*(*μ, σ*). Hence, the average model 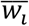 across cohorts was an unbiased estimator for the true coefficient, and therefore the squared penalty 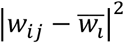 was incorporated to penalize the departure of each model j to the mean. The complete formulation was

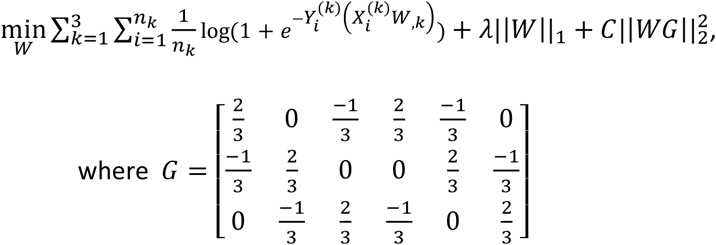

#### Computational speed of unsupervised dsMTL

Here, we analyzed the time efficiency in applying dsMTL_iNMF on two real datasets based on the real network latency. Two processed RNA-seq case-control cohorts comprising patients with schizophrenia (GSE164376^33^) and bipolar disorder (GSE134497^34^) were retrieved from the GEO database and converted into a matrix format for the analysis. As shown in **Supplementary Table 4**, the data were stored on servers in Mannheim and Heidelberg.

## Supporting information

supplementary results

## Acknowledgements

This study was supported by the Deutsche Forschungsgemeinschaft (DFG), SCHW 1768/1-1 and the German Federal Ministry of Education and Research (BMBF, grants 01KU1905A and 01ZX1904A).

## References

1. Jahanshad N, Kochunov PV, Sprooten E, et al. Multi-site genetic analysis of diffusion images and voxelwise heritability analysis: A pilot project of the ENIGMA–DTI working group. NeuroImage. 2013;81:455–469.

2. Schizophrenia Working Group of the Psychiatric Genomics C. Biological insights from 108 schizophrenia-associated genetic loci. Nature. 2014;511(7510):421–427.

3. Kochunov P, Jahanshad N, Sprooten E, et al. Multi-site study of additive genetic effects on fractional anisotropy of cerebral white matter: comparing meta and megaanalytical approaches for data pooling. NeuroImage. 2014;95:136–150.

4. Carter KW, Francis RW, Carter K, et al. ViPAR: a software platform for the Virtual Pooling and Analysis of Research Data. International journal of epidemiology. 2016;45(2):408–416.

5. Warnat-Herresthal S, Schultze H, Shastry KL, et al. Swarm Learning for decentralized and confidential clinical machine learning. Nature. 2021;594(7862):265–270.

6. Zhou J, Liu J, Narayan VA, Ye J, Alzheimer’s Disease Neuroimaging I. Modeling disease progression via multi-task learning. NeuroImage. 2013;78:233–248.

7. Gaye A, Marcon Y, Isaeva J, et al. DataSHIELD: taking the analysis to the data, not the data to the analysis. International journal of epidemiology. 2014;43(6):1929–1944.

8. Wilson RC, Butters OW, Avraam D, et al. DataSHIELD – New Directions and Dimensions. Data Science Journal. 2017;16.

9. Cao H, Zhou J, Schwarz E. RMTL: An R Library for Multi-Task Learning. Bioinformatics. 2018.

10. Yang Z, Michailidis G. A non-negative matrix factorization method for detecting modules in heterogeneous omics multi-modal data. Bioinformatics. 2016;32(1):1–8.

11. Xu Q, Xue H, Yang Q. Multi-platform gene-expression mining and marker gene analysis. International journal of data mining and bioinformatics. 2011;5(5):485–503.

12. Yuan H, Paskov I, Paskov H, Gonzalez AJ, Leslie CS. Multitask learning improves prediction of cancer drug sensitivity. Scientific reports. 2016;6:31619.

13. Cao H, Meyer-Lindenberg A, Schwarz E. Comparative Evaluation of Machine Learning Strategies for Analyzing Big Data in Psychiatry. International journal of molecular sciences. 2018;19(11).

14. Zhou J, Yuan L, Liu J, Ye J. A multi-task learning formulation for predicting disease progression. 2011:814.

15. Fujita N, Mizuarai S, Murakami K, Nakai K. Biomarker discovery by integrated joint non-negative matrix factorization and pathway signature analyses. Scientific reports. 2018;8(1):9743.

16. Tibshirani R. Regression shrinkage and selection via the lasso. Journal of the Royal Statistical Society Series B (Methodological). 1996:267–288.

17. Lichtenstein P, Yip BH, Björk C, et al. Common genetic determinants of schizophrenia and bipolar disorder in Swedish families: a population-based study. The Lancet. 2009;373(9659):234–239.

18. Matschinske J, Späth J, Nasirigerdeh R, et al. The FeatureCloud AI Store for Federated Learning in Biomedicine and Beyond. arXiv preprint arXiv:210505734. 2021.

19. Marcon Y, Bishop T, Avraam D, et al. Orchestrating privacy-protected big data analyses of data from different resources with R and DataSHIELD. PLoS computational biology. 2021;17(3):e1008880.

20. Lenz S, Hess M, Binder H. Deep generative models in DataSHIELD. BMC Med Res Methodol. 2021;21(1):64.

21. Zhang C, Liu J. Distributed Learning Systems with First-Order Methods. Foundations and Trends^®^ in Databases. 2020;9(1):1–100.

22. Xie L, Baytas IM, Lin K, Zhou J. Privacy-Preserving Distributed Multi-Task Learning with Asynchronous Updates. 2017:1195–1204.

23. Dahl M, Mancuso J, Dupis Y, et al. Private machine learning in tensorflow using secure computation. arXiv preprint arXiv:181008130. 2018.

24. Warnat-Herresthal S, Schultze H, Shastry KL, et al. Swarm Learning as a privacy-preserving machine learning approach for disease classification. 2020.

25. Friedman J, Hastie T, Tibshirani R. Regularization Paths for Generalized Linear Models via Coordinate Descent. Journal of Statistical Software. 2010;33(1).

26. Purcell S, Neale B, Todd-Brown K, et al. PLINK: a tool set for whole-genome association and population-based linkage analyses. American journal of human genetics. 2007;81(3):559–575.

27. Hu H, Salcic Z, Dobbie G, Zhang X. Membership Inference Attacks on Machine Learning: A Survey. arXiv preprint arXiv:210307853. 2021.

28. Fredrikson M, Lantz E, Jha S, Lin S, Page D, Ristenpart T. Privacy in pharmacogenetics: An end-to-end case study of personalized warfarin dosing. Paper presented at: 23rd {USENIX} Security Symposium ({USENIX} Security 14) 2014.

29. Lanz TA, Reinhart V, Sheehan MJ, et al. Postmortem transcriptional profiling reveals widespread increase in inflammation in schizophrenia: a comparison of prefrontal cortex, striatum, and hippocampus among matched tetrads of controls with subjects diagnosed with schizophrenia, bipolar or major depressive disorder. Translational psychiatry. 2019;9(1):151.

30. Tang B, Capitao C, Dean B, Thomas EA. Differential age- and disease-related effects on the expression of genes related to the arachidonic acid signaling pathway in schizophrenia. Psychiatry research. 2012;196(2-3):201–206.

31. Chen C, Cheng L, Grennan K, et al. Two gene co-expression modules differentiate psychotics and controls. Mol Psychiatry. 2013;18(12):1308–1314.

32. Fromer M, Roussos P, Sieberts SK, et al. Gene expression elucidates functional impact of polygenic risk for schizophrenia. Nature neuroscience. 2016;19(11):1442–1453.

33. A; K, R; K. GSE164376 dataset. https://www.ncbi.nlm.nih.gov/geo/query/acc.cgi?acc=GSE164376. Published 2021. Accessed.

34. Kathuria A, Lopez-Lengowski K, Vater M, McPhie D, Cohen BM, Karmacharya R. Transcriptome analysis and functional characterization of cerebral organoids in bipolar disorder. Genome medicine. 2020;12(1):34.

