## supplementary methods for "dsMTL - a computational framework for privacy-preserving, distributed multi-task machine learning"

#### Outline

- dsMTL framework
  - Distributed unsupervised method in dsMTL
  - Distributed supervised methods in dsMTL
- Introduction of DataSHIELD
- Data key mechanism
- Generate RNA-seq count data for case study 2
- Microarray expression Data pre-process

#### dsMTL framework

In dsMTL, we included four federated multi-task (FeMTL) and one machine learning (FeML) methods covering supervised and unsupervised learning procedures. All models followed the consistent formulation,

$$\min_{\theta} \mathcal{L}(\theta) + \lambda S(\theta) + C\aleph(\theta) \quad (1)$$

$\mathcal{L}(\theta)$ was the data fitting term (or loss function), the major determinant of the solutions of the model training. $\aleph(\theta)$ and $S(\theta)$ were the regularization/penalty terms with the aim to incorporate the prior information and prevent overfitting. $\aleph(\theta)$ was a non-smooth function for creating the sparsity, while $S(\theta)$ was smooth with the ability to stabilize the solution. $\lambda$ and $C$ were the hyper-parameters to control the strength of the penalty, $\lambda$ was learnt from cross-validation (CV) and $C$ was the constant.

There are three loss functions in dsMTL, achieving the tasks of regression, classification and matrix factorization. They are summarized in **Supplementary Table 1**.

| | Unsupervised Learning | Supervised Learning | |
|---|---|---|---|
| | Matrix factorization | Regression | Classification |
| Model | $[X_1, \dots, X_k, \dots X_t] = [(H + Hv_1) \times W_1, \dots, (H + Hv_t) \times W_t])$ | $f(x) = xw$ | $P(x) = \frac{1}{1 + e^{-(xw)}}$ |
| Loss function | $\min_{H, W_1 \dots, W_K, V_1 \dots, V_K} \sum_{k=1}^{t} \|\|X_k - (H + V_k)W_k\|\|_F^2$ | $\min_{w} \frac{1}{2n} \sum_{i=1}^{n} \|\|y_i - x_i w\|\|^2$ | $\min_{w} \frac{1}{n} \sum_{i=1}^{n} \log(1 + e^{-y_i(x_i w)})$ |
| Gradient | $\nabla_{H_{i,j}} = 2 \sum_{k=1}^{t} (H_{i,j} W_{k_i} W_{k_i}^T - X_{k_i} W_{k_i}^T)$ <br> $\nabla_{W_{k_{i,j}}} = 2 \sum_{m=1}^{n_t} (HV_k)_{m,i} \left[(HV_k)_{m,i} W_{k_{ij}} - X_{k_{m,j}}\right]$ <br> $\nabla_{V_{k_{i,j}}} = 2(V_{i,j} W_{k_i} W_{k_i}^T - X_{k_i} W_{k_i}^T)$ | $\nabla_w = \frac{1}{n}(x^T x w - x^T Y)$ | $\nabla_w = -\frac{1}{n} X^T \times \begin{bmatrix} \frac{y_1}{1 + e^{y_1(x_1 w)}} \\ \dots \\ \frac{y_n}{1 + e^{y_n(x_n w)}} \end{bmatrix}$ |

**Supplementary Table 1.** Summaries of loss functions used in dsMTL

***<u>Termination rules</u>*** Four termination rules were included in dsMTL to determine whether the optimization converges. The first three rules were applied to all methods in dsMTL, while the last was new designed for matrix factorization. The first rule checked whether the current objective value was close enough to 0. The second rule investigates the last two objective values and checks whether the decrement was close enough to 0. The third rule allowed the optimization to be performed for a certain maximum number of iterations. The last rule specific to matrix factorization was described in the next section.

#### Federated unsupervised method in dsMTL

To discover the hidden structure in heterogeneous, high-dimensional biological data, we integrated the integrative matrix factorization method[1] (iNMF) in our distributed learning framework, called dsMTL_iNMF. The major concept of dsMTL_iNMF is shown in **Supplementary Figure 1**, where the cohort matrices on three servers can be factorized simultaneously with the shared component matrix ($H$) and cohort-specific component matrices(V, W). The objective function was

$$\min_{\substack{H,\\ W_1\ldots,W_K,\\ V_1\ldots,V_K}} \sum_{k=1}^{t} ||X_k - (H + V_k)W_k||_F^2 + \lambda \sum_{k=1}^{t} ||V_k W_k||_F^2 + \lambda_s \sum_{k=1}^{t} |W_k|_1$$

The robustness of the model was due to the decoupled setting of $H$ and $V_k$, where $H$ was to capture the shared information across cohorts and $V_k$ was to capture the cohort-specific information. To integrate the more information into shared component H, the magnitude of cohort-specific component was penalized $\aleph(.) = \sum_{k=1}^{t} ||V_k W_k||_F^2$. The sparse term $S(.) = \sum_{k=1}^{t} |W_k|_1$ was used to remove the redundant coefficients from the component matrices.

##### Distributed variables update

$$W_{k_{ij}} \leftarrow W_{ij} \frac{((H+V_k)^T X_k)_{i,j}}{((H^T H + H V_k{}^T + H^T V_k + (1+\lambda) V_k{}^T V_k) W_k)_{i,j} + \lambda_s} \quad (2)$$

$$V_{k_{ij}} \leftarrow V_{ij} \frac{(X_k W_k^T)_{i,j}}{(H W_k W_k^T + (1+\lambda) V_k W_k W_k^T)_{i,j} + \lambda_s} \quad (3)$$

$$H_{ij} \leftarrow H_{ij} \left( \frac{X_1 W_1^T + \cdots + X_t W_t^T}{(H+V_1) W_1 W_1^T + \cdots + (H+V_t) W_t W_t^T} \right)_{i,j} \quad (4)$$

The variables were updated for non-federated applications as demonstrated in formulas (2) to (4). In the federated scenario, the cohort-specific variables $W_k$ and $V_k$ were updated on server k using the local data as formulas (2) and (3). The shared matrix $H$ was updated on the client after receiving summary data (see **Supplementary Figure 1**) from all servers, where these were not-disclosed and calculated behind a given institution's firewall. The distributed update of $H$ was

$$H_{ij} \leftarrow H_{ij} \left( \frac{server_1(X_1 W_1^T) + \cdots + server_t(X_t W_t^T)}{server_1((H+V_1) W_1 W_1^T) + \cdots + server_t((H+V_t) W_t W_t^T)} \right)_{i,j} \quad (5)$$

After the aggregation, the client updates $H$ and a new iteration begin. The communications between the client and the servers are illustrated in **Supplementary Figure 1**.

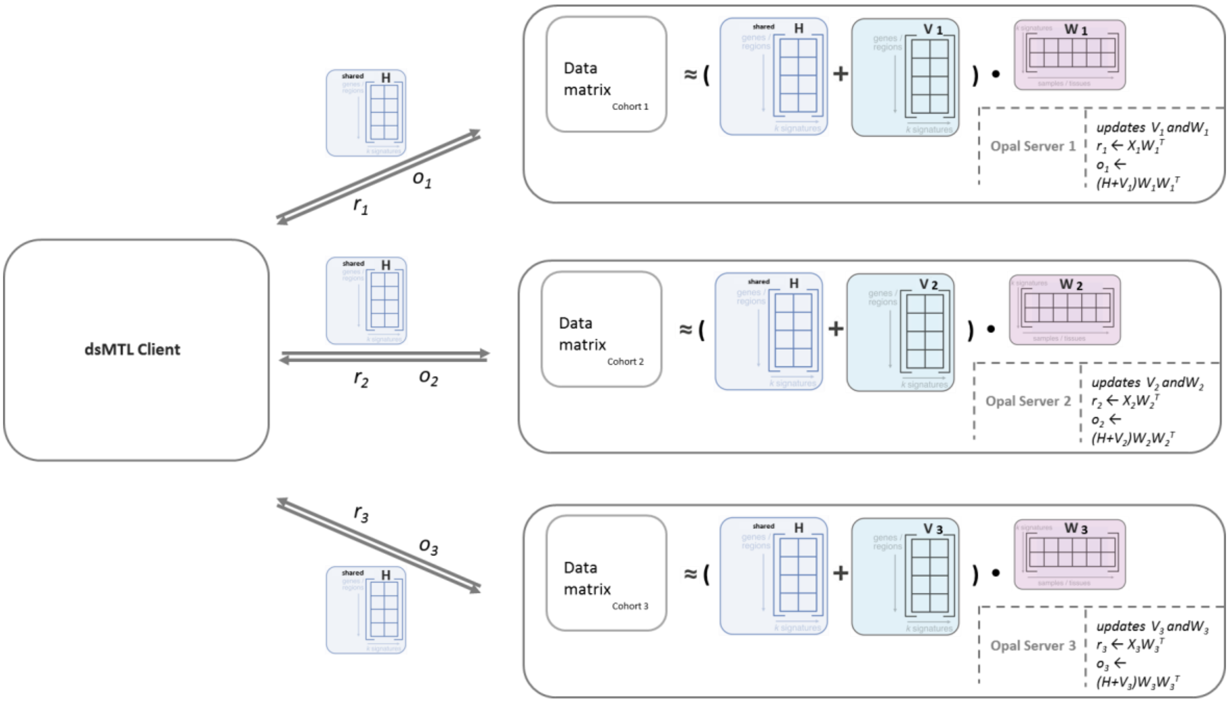


**Supplementary Figure 1**. Communications between the client and opal servers for dsMTL_iNMF

#### Algorithms

##### Distributed solver

For the privacy-preserving purpose, only the shared matrix $H$ was returned. The distributed solver of dsMTL_iNMF is shown in Algorithm 1.

| **Algorithm 1**, Solver of distributed iNMF in dsMTL |
| --- |
| **Input**: $\lambda > 0$, $\lambda_s > 0$, maxIter $> 0$, $H$, $W_1 \dots, W_K$, $V \dots, V_K$ |
| **Output**: $H$ |
| 1: **for** $i = 1$ to maxIter **do** |
| 2: Update $H$ according to (5) on client |
| 3: Send $H$ to all servers |
| 3: Update $W_1 \dots, W_K$ according to (2) on server $1, \dots, k$ |
| 4: Update $V_1 \dots, V_K$ according to (3) on server $1, \dots, k$ |
| 5: Send summary statistics back client according to (5) |
| 6: If termination rule satisfied, **return** |

##### Termination rules

For dsMTL_iNMF, we provided an additional termination rule developed in the ButchR package[2] to determine the convergence of the algorithm. In this method, each of the samples was assigned to a hidden factor (clustering membership) by $\mathrm{j} = \arg\max_j |H_{i,j}|$ at every iteration. The convergence was determined when the assignments of samples remained unchanged. By default, if the samples were assigned to the same hidden factors consistently for over 10 iterations, the memberships were seen as stable, and the algorithm stopped. The rationale behind this procedure is that in order to maximize the power of clustering, the variance of the determined memberships must be small. Therefore, the proposed rule terminates the algorithm when the samples found stable memberships, such that the clustering can make a stable decision.

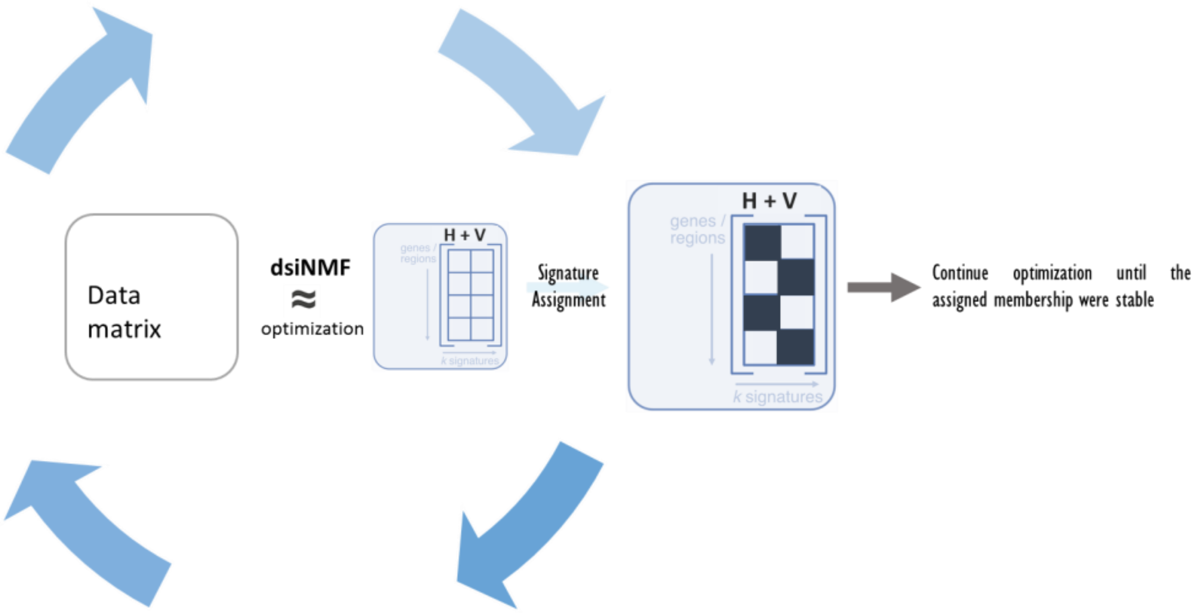


**Supplementary Figure 2**. Schematic illustration of cluster membership optimization.

**Distributed model training**

In the default setting, Algorithm 1 was performed with 10 random initial points to approximately sample the distribution of the local optima considering the non-convex nature of the problem. The initialization of these component matrices were uniformly sampled from $[0, 2]$. For each initialization, Algorithm 1 was performed. A set of shared matrices were returned as the final results for subsequent analysis

**Algorithm 2** Training procedure of iNMF in dsMTL

**Input**: $\lambda > 0$, $\lambda_s > 0$, maxIter $> 0$, rank, nInitialization, $\{X_1, \dots X_k, \dots X_t\}$

**Output**: $H_1$, $H_2, \dots$

1: **for** $i = 1$ to nInitialization **do**

2: Initialize $H_i \sim U_{n \times rank}(0,1)$, for each k, $V_k \sim U_{n \times rank}(0,1)$, $W_k \sim U_{rank \times p_k}(0,1)$

3: $H_i$ =**Algorithm 1** ($\lambda = \lambda, \lambda_s = \lambda_s, H = H_i, \{W_1, \dots, W_t\}, \{V_1, \dots, V_t\}$)

4: **end for**

#### Federated supervised methods in dsMTL

We included one machine-learning (ML) and three multi-task learning (MTL) algorithms into supervised methods of dsMTL. In the federated scenario, the ML model was trained by averaging the summary statistics from geo-distributed cohorts with the synchronous communication, which leads to a model equivalent to the standalone ML model training on the concatenated cohorts. MTL, in the federated scenario, exchanges a small amount of information by regularization, such that the commonality of multi-cohort models was reinforced, but the cohort-specific element remained unchanged. In dsMTL, dsLasso was included as the federated ML variant of Lasso[3]. The federated multi-task learning methods were adapted from our previously published package RMTL[4]. These methods adopted various strategies of cross-cohort regularization to select joint features, explore the low-rank structure and incorporate the network structure in multiple tasks.

In the next sections, we introduce the theoretical derivations of each method to form the federated algorithm. Then a federated optimization framework is derived that was applied to all supervised methods. At the end, we show an executable algorithms for solving the objective and training the model.

#### Models

For each method, the objective function is first introduced as the major problem to solve. Second, we derive the subgradient of the objective to characterize the properties of the optima. Third, since all models are sparse, we aimed to solve the entire regularization tree[5] with a given positive $\lambda$ sequence in decreasing order. The $\lambda_{max}$ , the estimate of largest $\lambda$ in the $\lambda$ sequence, was derived from the subgradient, such that $\lambda_{max}$ was the smallest $\lambda$ guaranteeing the existence of 0 optima. Last, to solve the non-smooth objective efficiently, the proximal mapping was applied, and the proximal point estimator was derived as the solution of the iteration-level sub-problem (see next section).

***Lasso (dsLasso)***

Objective function:

$$\min_{w} \frac{1}{n}\sum_{i=1}^{n} \mathcal{L}(w; X, Y) + \lambda|w| + C||w||_2^2$$

Subgradient:

$$\partial_w = \nabla_w \mathcal{L} + 2Cw + \lambda \frac{w}{|w|}$$

Estimated $\lambda_{max}$:

$$\lambda_{max} = \begin{cases} \frac{1}{n}\max_j |X_{,j}^T Y| & for\ least\ square\ loss \\ \frac{1}{2n}\max_j |X_{,j}^T Y| & for\ logit\ loss \end{cases}$$

Proximal point estimation:

$$\operatorname{prox}_{f=\frac{\lambda|x|}{L}}(w) = \operatorname{sign}(w)\max\{w-\lambda, 0\}$$

The Lasso model[3] aimed to learn a sparse parameter vector $w$. $\lambda$ was identified by cross-validation. $||w||_2^2$ was used to stabilize the solution and incorporate the correlated features. $C$ was selected by the user.

***MTL with Feature selection (dsMTL_L21)***

Objective function

$$\min_W \sum_{k=1}^{t} \sum_{i=1}^{n_k} \frac{1}{n_k} \mathcal{L}\left(W_{,k}; X_i^{(k)}, Y_i^{(k)}\right) + \lambda \sum_{j=1}^{p} \sqrt{||W_{j,}||_2^2} + C||W||_2^2$$

Subgradient:

$$\partial_{W_{j,}} = \nabla_{W_{j,}} \mathcal{L} + 2CW_{j,} + \lambda v,\ v = \{x \in R^t: ||x||_2 \leq 1\}$$

Estimated $\lambda_{max}$:

$$\lambda_{max} = \begin{cases} \max_j \sqrt{\sum_{k=1}^{t} \left(\frac{< X_{,j}^{(k)}, Y^{(k)} >}{n_k}\right)^2} & for\ least\ square\ loss \\ \max_j \sqrt{\sum_{k=1}^{t} \left(\frac{< X_{,j}^{(k)}, Y^{(k)} >}{2n_k}\right)^2} & for\ logit\ loss \end{cases}$$

Proximal point estimation:

$$\operatorname{prox}_{f=\frac{\lambda||x||_2}{L}}(w) = \left(1 - \frac{\lambda}{\max\left\{\left||w|\right|_2, \lambda\right\}}\right) w$$

The method[6] aimed to find a model with the same set of features. For this, $\sum_{j=1}^{p} \sqrt{||W_{j,}||_2^2}$ was used to penalize the magnitudes of the coefficients of a given feature across the datasets. The $W = p \times t$ was the solution matrix of $t$ tasks and p features.

***<u>MTL with low-rank structure</u>(dsMTL_Trace)***

Objective function:

$$\min_W \sum_{k=1}^{t} \sum_{i=1}^{n_k} \frac{1}{n_k} \mathcal{L}\left(W_{,k}; X_i^{(k)}, Y_i^{(k)}\right) + +\lambda||W||_* + C||W||_2^2$$

Subgradient:

$$\partial_W = \nabla_W \mathcal{L} + 2CW + \lambda \partial ||W||_*$$

Estimated $\lambda_{max}$:

$$\lambda_{max} = \begin{cases} \max_j \sigma_1(\left[\frac{X^{(1)^T} Y^{(1)}}{n_1}, \dots, \frac{X^{(t)^T} Y^{(t)}}{n_t}\right]) & for\ least\ square\ loss \\ \max_j \sigma_1(\left[\frac{X^{(1)^T} Y^{(1)}}{2n_1}, \dots, \frac{X^{(t)^T} Y^{(t)}}{2n_t}\right]) & for\ logit\ loss \end{cases}$$

Where $\sigma_1(A)$is the largest singular value of matrix A

Proximal point estimation:

$$\operatorname*{prox}_{f=\frac{\lambda||x||_*}{L}}(W) = U \times I_{\max\{\sigma-\lambda,0\}} \times V,$$

where $W = U\Sigma V$, $\sigma$ is the diagonal vector of $\Sigma$

The method[7] aimed to identify the coefficient vectors of multiple cohorts existing in the compressed low-dimensional space. For this, the trace norm of the coefficient matrix was used to compress the models' space.

***MTL with network structure(dsMTL_Net)***

Objective function

$$\min_W \sum_{k=1}^{t}\sum_{i=1}^{n_k}\frac{1}{n_k}\mathcal{L}\left(W_{,k}; X_i^{(k)}, Y_i^{(k)}\right) + \lambda||W||_1 + C||GW||_2^2$$

Subgradient:

$$\partial_W = \nabla_W \mathcal{L} + 2CGG^T + \lambda\frac{W}{|W|}$$

Estimated $\lambda_{max}$:

$$\lambda_{max} = \begin{cases} \max\limits_{j,k}\frac{{X_{,j}^{(k)}}^T Y^{(k)}}{n_k} & for\ least\ square\ loss \\ \max\limits_{j,k}\frac{{X_{,j}^{(k)}}^T Y^{(k)}}{2n_k} & for\ logit\ loss \end{cases}$$

Where $\sigma_1(A)$is the largest singular value of matrix A

Proximal point estimation:

$$\operatorname*{prox}_{f=\frac{\lambda|x|}{L}}(W) = \mathrm{sign(W)}\max\{|W| - \lambda, 0\}$$

The method aimed to incorporate the relationships between cohorts as a graph into the model training procedure. $||GW||_2^2$ was used for this aim, where G was an pre-defined matrix describing the network structure. More details of G for variant applications can be found in [8]. $||W||_1$ was used to remove redundant coefficients. $\lambda$ was identified by cross-validation.

#### Distributed Optimization procedure

To solve these composite objective functions efficiently in the same framework, we rewrite the objective (1) as

$$\min_x F(x) + \lambda\Omega(x) \qquad (9)$$

where $F(x) = \mathcal{L}(\theta) + CS(\theta)$ was smooth component function and $\Omega(x) = \aleph(\theta)$ was non-smooth

**Solving sub-problem in each iteration**

Given the Lipschitz constant $L$ of the objective function above, the sequence of estimation points $\{x_0, x_1, x_2, \dots\}$ were found by solving the below iteration-wise sub-problem (9)

$$x_{i+1} = \arg\min_{y} \mathcal{M}_{L,x_i}(y) \quad (10)$$

$$\mathcal{M}_{L,x_i}(y) = F(x_i) + \langle \nabla F(x_i), y - x_i \rangle + \frac{L}{2}\left|\left|y - x_i\right|\right|_2^2 + \lambda\Omega(x) \quad (11)$$

The first three terms on the right side were the second order approximation of $F(.)$ using Tylor expansion on point $x_i$. After re-organization, we have (12) equal to (10).

$$x_{i+1} = \arg\min_{y} \frac{L}{2}\left(y - \left(x_i - \frac{\nabla F(x_i)}{L}\right)\right)^2 + \lambda\Omega(x) \quad (12)$$

Since $x_i - \frac{\nabla F(x_i)}{L}$ was known after the $i$th iterations, the above problem was applicable to the proximal algorithm framework[10]. For all sparse regularizations ($\Omega(x)$) used in dsMTL, they can be simplified and solved analytically in (13), and the results were derived and summarized above (see the "Proximal point estimation") for each dsMTL method.

$$x_{i+1} = \underset{f=\frac{\lambda\Omega(x)}{L}}{\operatorname{prox}} \left(x_i - \frac{\nabla F(x_i)}{L}\right) \quad (13)$$

#### Line search

Since $L$ was unknown in our framework, we estimated it using a backtracking line search approach. Set an increasing sequence of $L \in \{L_0, 2L_0, 4L_0, 16L_0, \dots\}$given an constant $L_0$, the smallest $L$ satisfying the condition $\mathcal{M}_{L,x_i}(x_{i+1}) \geq F(x_{i+1})$ was selected. Here, $x_{i+1}$ was determined based on (12).

#### Federated computation

For supervised MTL, the variable matrix $\boldsymbol{W} = \boldsymbol{p} \times \boldsymbol{t} = [\boldsymbol{w}_{,1}, \boldsymbol{w}_{,2}, \dots, \boldsymbol{w}_{,t}]$, where each column represents one task. So distributed proximal operator was:

$$W_{i+1} = \underset{f=\frac{\lambda\Omega(x)}{L}}{\operatorname{prox}} \left(W_i - \frac{\nabla F(W_i)}{L}\right) = \underset{f=\frac{\lambda\Omega(x)}{L}}{\operatorname{dist\,prox}} \left(\left[w_{,1_i} - \frac{\nabla F\left(w_{,1_i}; D^{(1)}\right)}{L}, \quad \dots, \quad w_{,t_i} - \frac{\nabla F\left(w_{,t_i}; D^{(t)}\right)}{L}\right]\right) \quad (14)$$

where $w_{,k_i} - \frac{\nabla F\left(w_{,k_i}\right)}{L}$ was calculated on server k and sent back. $\{D_{1,\dots}, D_t\}$ represented the data on t servers. Similarly, objective function $O(W)$ has to be evaluated in a distributed fashion,

$$O(W) = \operatorname{dist} O(W) = \sum_{k=1}^{t} F\left(W_{,k}; D^{(k)}\right) + \lambda\Omega(W)$$

For supervised ML, the information aggregation was different. The variable vector $\boldsymbol{w} = \boldsymbol{p} \times \boldsymbol{1}$. The distributed proximal operator is

$$w_{i+1} = \underset{f=\frac{\lambda\Omega(x)}{L}}{\operatorname{prox}} \left(w_i - \frac{\nabla F(w_i)}{L}\right) = \underset{f=\frac{\lambda\Omega(x)}{L}}{\operatorname{dist\,prox}} \left(w_i - \frac{1}{L}\left(\sum_{j=1}^{t} \nabla\mathcal{L}\left(w_i; D^{(j)}\right)\frac{n_t}{n} + C\nabla\aleph(w_i)\right)\right)$$

where $\nabla\mathcal{L}(w_i; D^{(j)})$ was calculated on server j and sent back. Similarly, objective function $O(w)$ has to be evaluated in distributed federated fashion,

$$O(w) = \mathrm{dist}\, O(w) = \sum_{j=1}^{t} \mathcal{L}(w_i; D^{(j)}) \frac{n_j}{n} + C\aleph(w_i) + \lambda\Omega(w).$$

#### Accelerated algorithms

##### ***Distributed solver***

To accelerate the optimization procedure, we applied Nesterov's acceleration approach[9,11,12]. In the beginning of iteration $i$, the search point was first defined as the weighted combination of the results from the previous two steps: $\mathrm{S_i} = \frac{\alpha_{i-1}}{\alpha_i}\mathrm{x_i} + \frac{1-\alpha_{i-1}}{\alpha_i}\mathrm{x_{i-1}}$. Then the formulas (13) was applied on the $\mathrm{S_i}$.

**Algorithm 3** Distributed solver of supervised learning methods in dsMTL

**Input**: $\lambda > 0$, $L_0 > 0$, $W_0$, $maxIter > 0$

**Output**: $W_{i+1}$

Initialize $W_1 = W_0$, $\alpha_{-1} = \alpha_0 = 0$, and $L = L_0$

**for** $i = 1$ to $maxIter$ **do**

Set $\mathrm{S_i} = \mathrm{W_i} + \frac{\alpha_{i-1}-1}{\alpha_i}(\mathrm{W_i} - \mathrm{W_{i-1}})$

Find smallest $L \in \{L_{i-1}, 2L_{i-1}, 4L_{i-1}, 16L_{i-1}, \ldots\}$ such that

$$\mathcal{M}_{L,x_i}(W_{i+1}) \geq \mathrm{dist}\, O(W_{i+1}),$$

where $W_{i+1} = \mathrm{dist}\ \underset{f=\frac{\lambda\Omega(x)}{L}}{\mathrm{prox}} \left(W_i - \frac{\nabla F(W_i)}{L}\right)$

Set $L_i = L$, and $\alpha_{i+1} = \frac{1+\sqrt{1+4\alpha_i^2}}{2}$

If termination rule satisfied, **return**

**end for**

##### ***Training for sparse model***

In the high-dimensional data analysis, the performance of sparse ML models was highly related to the accuracy of sparse structure identification, thus $\lambda$ selection was crucial. In dsMTL, we trained the entire

**Algorithm 4** Training procedure of sparse models in dsMTL

**Input**: $\lambda_1 > \lambda_2 > \cdots > 0$

**Output**: $W_1, W_2, \ldots$

Initialize $W_0 = p \times t = 0$

**for** $i = \{1, 2, \ldots\}$ **do**

$W_i =$ **Algorithm 3** ($\lambda = \lambda_i, L_0 = 1, W_0 = W_{i-1}, maxIter = 100$)

**end for**

regularization tree for a given hyper-paratemer C. Similar to the study[13], we estimated the $\lambda_{max}$ as the largest $\lambda$ of the sequence from the data. $\lambda_{max}$ was selected by looking for the smallest $\lambda$ such that the equation $\partial_W\big(F(x)+\lambda\Omega(x)\big)\ni \mathbf{0}$ hold. Due to the differential objective functions, $\lambda_{max}$ of calssification model was different from that of regression model. The $\lambda_{min}$ was determined based on $\lambda_{max}$, i.e. $0.1\lambda_{max}$. Then the entire sequence was interpolated based on the log scale of $\lambda_{max}$ and $\lambda_{min}$. For each method, $\lambda_{max}$ was therotically different, and summarized above.

***Cross-validation***

We set up cross-cohort and in-cohort CV in dsMTL for all ML/MTL methods. For cross-cohort CV, t folds CV were established for *t* cohorts. In fold *i*, cohort *i* was as the test cohort and the model was trained on rest cohorts. The prediction performances were averaged and used to select $\lambda$. Such CV aimed to identify a $\lambda$ with an optimized generalization performance. For k-folds in-cohort CV, the samples of each cohort were randomly separated into k folds, such that the test folds across cohorts were combined for testing, and the training folds were combined for training. Such CV aimed to identify a $\lambda$ with the most representative sparse model across all cohorts.

#### Introduction of DataSHIELD

DataSHIELD[14] is a platform software supporting federated data analysis without disclosing personally identifiable information. Two modules were included, the Ranalytic environment and the data warehouse opal. To mitigate the risk of sensitive data disclosure, the design of DataSHIELD considers to broader aspects: software architecture and statistics. The architecture of DataSHIELD provides several non-disclosure mechanisms to improve the system security, such as 1) data analysis only occurs behind the firewall; 2) each server is only allowed to communicate with a single client of a fixed IP; 3) the network communication was protected by the SSL protocol; 4) an R parser was implemented on the DataSHIELD server to reject unpermitted behaviors. A comprehensive set of permission settings were provided for data providers to fully control access to their data. These permissions were about users, data and functions for characterizing i.e. "*which users* could perform *what behaviors* to *what data"*. In an extremely secure example, a user could be granted to access a dataset but cannot perform any actions because no functions were granted. With these settings, DataSHIELD allows to customize the security strategies according to the specific requirement of the applications. From a statistical perspective, DataSHIELD assumes sharing summary statistics are safe for privacy-preserving applications. Such assumption is quite common in the biomedical field, and there is a large number of websites providing summary data for free download, such as the GWAS summary data of certain traits[15] and eQTLs of tissues[16]. Another study[17] confirmed the non-disclosure property of DataSHIELD for regression analysis from a biostatistical perspective.

### Data key mechanism

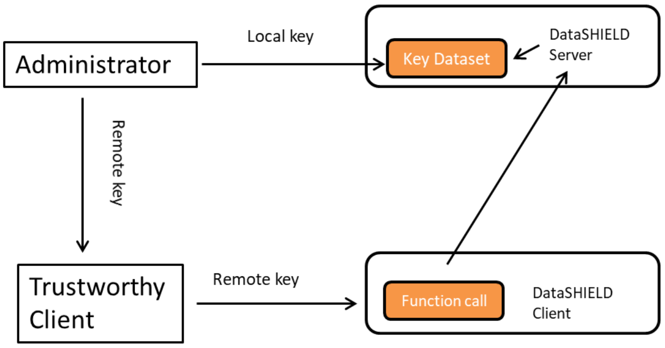


**Supplementary Figure 3**. Schematic illustration of the data key mechanism

This mechanism allows the authorized users to obtain the complete model identified by multi-task learning from the server. The administrator generates two keys, stores the local key in the key database, and gives the remote key to a trustworthy user. Then by sending the remote key, the client is seen as the data provider of the server, and can retrieve the complete model from the multi-task learning method.

This mechanism was built for two reasons. 1) The custom-defined functions in DataSHIELD cannot obtain identity information from the users, and 2) specifying the identity of users via the IP address is not sufficiently safe.

### Generate RNA-seq count data for case study 2

The RNA-seq count data was generated using the Negative Binomial distribution (NB distribution), which was the most common distribution used to model RNA-seq data. In case study 2, a two-cohort scenario was simulated. Four tests were conducted for different severity of heterogeneity. Here the severity of heterogeneity was characterized by the proportion (20%, 40%, 60% and 80%) of genes in the shared signature over all diagnosis-associated genes. The simulation procedure contained the following steps. First, the background data of two cohorts $X_1$and $X_2$were generated as sampled from the NB distribution $X_1 \sim \mathrm{NB}_{p \times n_1 \times}(\mathrm{r}=2, \mathrm{p}=0.3)$ and $X_2 \sim \mathrm{NB}_{p \times n_2}(\mathrm{r}=2, \mathrm{p}=0.3)$, where $p$ was shared gene dimension, $n_1$and $n_2$ referred to the respective sample size. Then in cohort $i$, the first 50% samples, $X_i[,1{:}\frac{n_i}{2}]$, were selected as patients while the first 50% genes, $X_i[1{:}\frac{p}{2},]$, were selected as the diagnosis-related genes. According to the specific proportion $\varphi\epsilon\{20\%, 40\%, 60\%, 80\%\}$ of shared genes over all signature genes, the shared and cohort-specific disease effect was added to the background data of patients. Specifically, the diagnosis-related effect shared by both cohorts was added as $X_i\left[1{:}\frac{p\varphi}{2}, 1{:}\frac{n_i}{2}\right] = X_i\left[1{:}\frac{p\varphi}{2}, 1{:}\frac{n_i}{2}\right] + \mathrm{NB}_{\frac{p\varphi}{2} \times \frac{n_i}{2}}(\mathrm{r}=2, \mathrm{p}=0.002)$ for cohort$i$. The diagnosis-related effect specific to cohort 1 was added as $X_1\left[\frac{2+p\varphi}{2}{:}\frac{p(1+\varphi)}{4}, 1{:}\frac{n_1}{2}\right] = X_1\left[\frac{2+p\varphi}{2}{:}\frac{p(1+\varphi)}{4}, 1{:}\frac{n_1}{2}\right] +$

$\mathrm{NB}_{\frac{p(1-\varphi)}{4}\times\frac{n_1}{2}}(\mathrm{r}=2,\mathrm{p}=0.002)$. The diagnosis-related effect specific to cohort 2 was added as $X_2\left[\frac{4+p(1+\varphi)}{4}:\frac{p}{2},1:\frac{n_2}{2}\right]=X_1\left[\frac{4+p(1+\varphi)}{4}:\frac{p}{2},1:\frac{n_2}{2}\right]+\mathrm{NB}_{\frac{p(1-\varphi)}{4}\times\frac{n_2}{2}}(\mathrm{r}=2,\mathrm{p}=0.002)$. Here the specific effects were not overlapped between cohorts 1 and 2.

#### Microarray expression data pre-processing

Four independent cortical microarray gene expression datasets from schizophrenia case-control cohorts were used in this study. Three datasets were downloaded from the GEO repository with id: GSE53987, GSE21138 and GSE35977. A detailed data description can be found on GEO and the respective original studies[18-20]. The fourth dataset was the HBCC microarray dataset (dbGAP ID: phs000979.v3.p2). Th data description and sample acquisition methods can be found on dbGAP and the original publication[21].

For GSE53987 and GSE21138, the expression levels were measured using the Affymetrix GeneChip Human Genome U133 Plus 2.0 Array, while the data of GSE35977 was measured using Affymetrix Human Gene 1.0 ST Array. A consistent pre-processing procedure was applied to all datasets. First, the raw data was extracted by the function *ReadAffy*() of the R package affy 1.64.0[22], followed by the rma[23] (Robust Multi-array Average) procedure for normalization. Values from multiple probes related to the same gene were averaged. Second, subjects with ages < 18 were excluded. Third, outliers were deleted as those outside of four standard deviations from the mean of the first two principal components. Fourth, 10 surrogate variables were determined using SVA[24] from the R package sva 3.34.0. Fifth, multiple linear regression analysis was used to correct for the effect of potential confounders with the covariates age, age$^2$, sex, PMI, pH, RIN, batch ID and 10 surrogate variables. Sixth, the resulting expression genes were z-standardized.

HBCC data was normalized and quality controlled as previously described[21]. First, we extracted the raw dlpfc expression data using the function *read.idat*() from the R package limma 3.42.2 [25]. Second, we corrected for background noise using the negative probes followed by quantile normalization and log-transformation. Third, we retained the robustly expressed probes as those with a detection p-value<0.01 in at least half of individuals. Prior to sva analysis, the missing “pH” and “PMI” values were imputed using the average of available data. The covariates contained age, age$^2$, sex, PMI, pH, RIN, ethnicity and 10 surrogate variables. The cohort contained 321 healthy controls and 191 patients with schizophrenia. All four datasets shared 8013 overlapping genes.

**References**


1. Yang Z, Michailidis G. A non-negative matrix factorization method for detecting modules in heterogeneous omics multi-modal data. *Bioinformatics.* 2016;32(1):1-8.
2. Quintero A, Hubschmann D, Kurzawa N, et al. ShinyButchR: Interactive NMF-based decomposition workflow of genome-scale datasets. *Biology methods & protocols.* 2020;5(1):bpaa022.
3. Tibshirani R. Regression shrinkage and selection via the lasso: a retrospective. *Journal of the Royal Statistical Society: Series B (Statistical Methodology).* 2011;73(3):273-282.
4. Cao H, Zhou J, Schwarz E. RMTL: An R Library for Multi-Task Learning. *Bioinformatics.* 2018.
5. Zou H, Hastie T. Regularization and variable selection via the elastic net. *Journal of the Royal Statistical Society: Series B (Statistical Methodology).* 2005;67(2):301-320.
6. Liu J, Ji S, Ye J. Multi-task feature learning via efficient l2, 1-norm minimization. Paper presented at: Proceedings of the Twenty-Fifth Conference on Uncertainty in Artificial Intelligence2009.
7. Pong TK, Tseng P, Ji S, Ye J. Trace Norm Regularization: Reformulations, Algorithms, and Multi-Task Learning. *SIAM Journal on Optimization.* 2010;20(6):3465-3489.
8. Cao H, Schwarz E. An Tutorial for Regularized Multi-task Learning using the package RMTL. The Comprehensive R Archive Network. Accessed.
9. Beck A, Teboulle M. A fast iterative shrinkage-thresholding algorithm for linear inverse problems. *SIAM journal on imaging sciences.* 2009;2(1):183-202.
10. Parikh N, Boyd S. Proximal algorithms. *Foundations and Trends® in Optimization.* 2014;1(3):127-239.
11. Nesterov Y. Gradient methods for minimizing composite functions. *Mathematical Programming.* 2012;140(1):125-161.
12. Liu J, Jieping Y. Efficient L1/Lq Norm Regularization.
13. Friedman J, Hastie T, Tibshirani R. Regularization Paths for Generalized Linear Models via Coordinate Descent. *Journal of Statistical Software.* 2010;33(1).
14. Wilson RC, Butters OW, Avraam D, et al. DataSHIELD – New Directions and Dimensions. *Data Science Journal.* 2017;16.
15. Zheng J, Erzurumluoglu AM, Elsworth BL, et al. LD Hub: a centralized database and web interface to perform LD score regression that maximizes the potential of summary level GWAS data for SNP heritability and genetic correlation analysis. *Bioinformatics.* 2017;33(2):272-279.
16. Consortium GT. Human genomics. The Genotype-Tissue Expression (GTEx) pilot analysis: multitissue gene regulation in humans. *Science.* 2015;348(6235):648-660.
17. Jones EM, Sheehan NA, Masca N, Wallace SE, Murtagh MJ, Burton PR. DataSHIELD – shared individual-level analysis without sharing the data: a biostatistical perspective. *Norsk Epidemiologi.* 2012;21(2).
18. Lanz TA, Reinhart V, Sheehan MJ, et al. Postmortem transcriptional profiling reveals widespread increase in inflammation in schizophrenia: a comparison of prefrontal cortex, striatum, and hippocampus among matched tetrads of controls with subjects diagnosed with schizophrenia, bipolar or major depressive disorder. *Translational psychiatry.* 2019;9(1):151.
19. Tang B, Capitao C, Dean B, Thomas EA. Differential age- and disease-related effects on the expression of genes related to the arachidonic acid signaling pathway in schizophrenia. *Psychiatry research.* 2012;196(2-3):201-206.
20. Chen C, Cheng L, Grennan K, et al. Two gene co-expression modules differentiate psychotics and controls. *Molecular psychiatry.* 2013;18(12):1308-1314.

21. Fromer M, Roussos P, Sieberts SK, et al. Gene expression elucidates functional impact of polygenic risk for schizophrenia. *Nature neuroscience.* 2016;19(11):1442-1453.
22. Gautier L, Cope L, Bolstad BM, Irizarry RA. affy--analysis of Affymetrix GeneChip data at the probe level. *Bioinformatics.* 2004;20(3):307-315.
23. Bolstad BM, Irizarry RA, Astrand M, Speed TP. A comparison of normalization methods for high density oligonucleotide array data based on variance and bias. *Bioinformatics.* 2003;19(2):185-193.
24. Leek JT, Johnson WE, Parker HS, Jaffe AE, Storey JD. The sva package for removing batch effects and other unwanted variation in high-throughput experiments. *Bioinformatics.* 2012;28(6):882-883.
25. Ritchie ME, Phipson B, Wu D, et al. limma powers differential expression analyses for RNA-sequencing and microarray studies. *Nucleic acids research.* 2015;43(7):e47-e47.
