## supplementary results for "dsMTL - a computational framework for privacy-preserving, distributed multi-task machine learning"

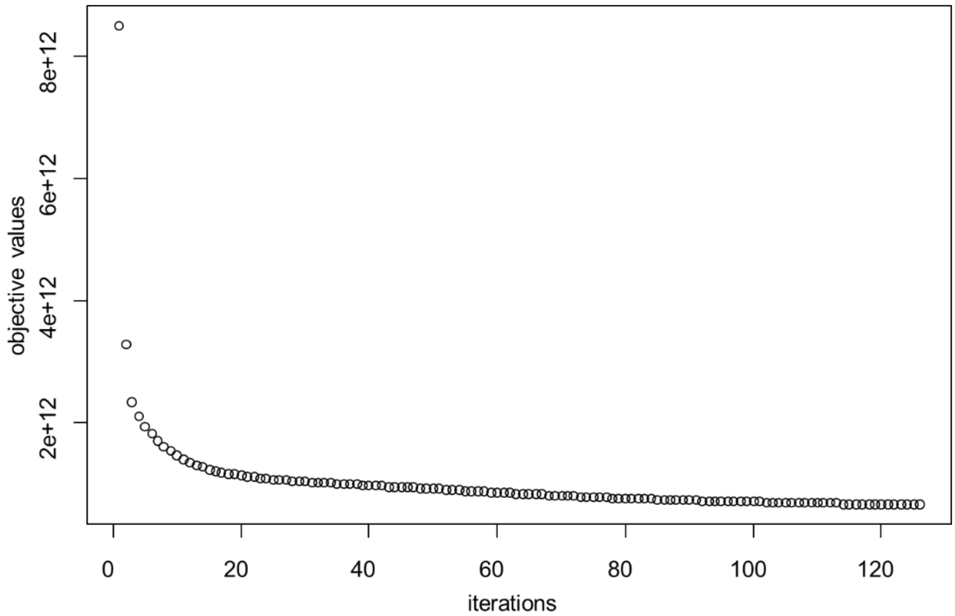


**Supplementary Figure 1**: the curve of objectives training dsMTL_iNMF with an initialization in case study 5. 100 iterations were sufficient to converge to a solution with an acceptable precision.

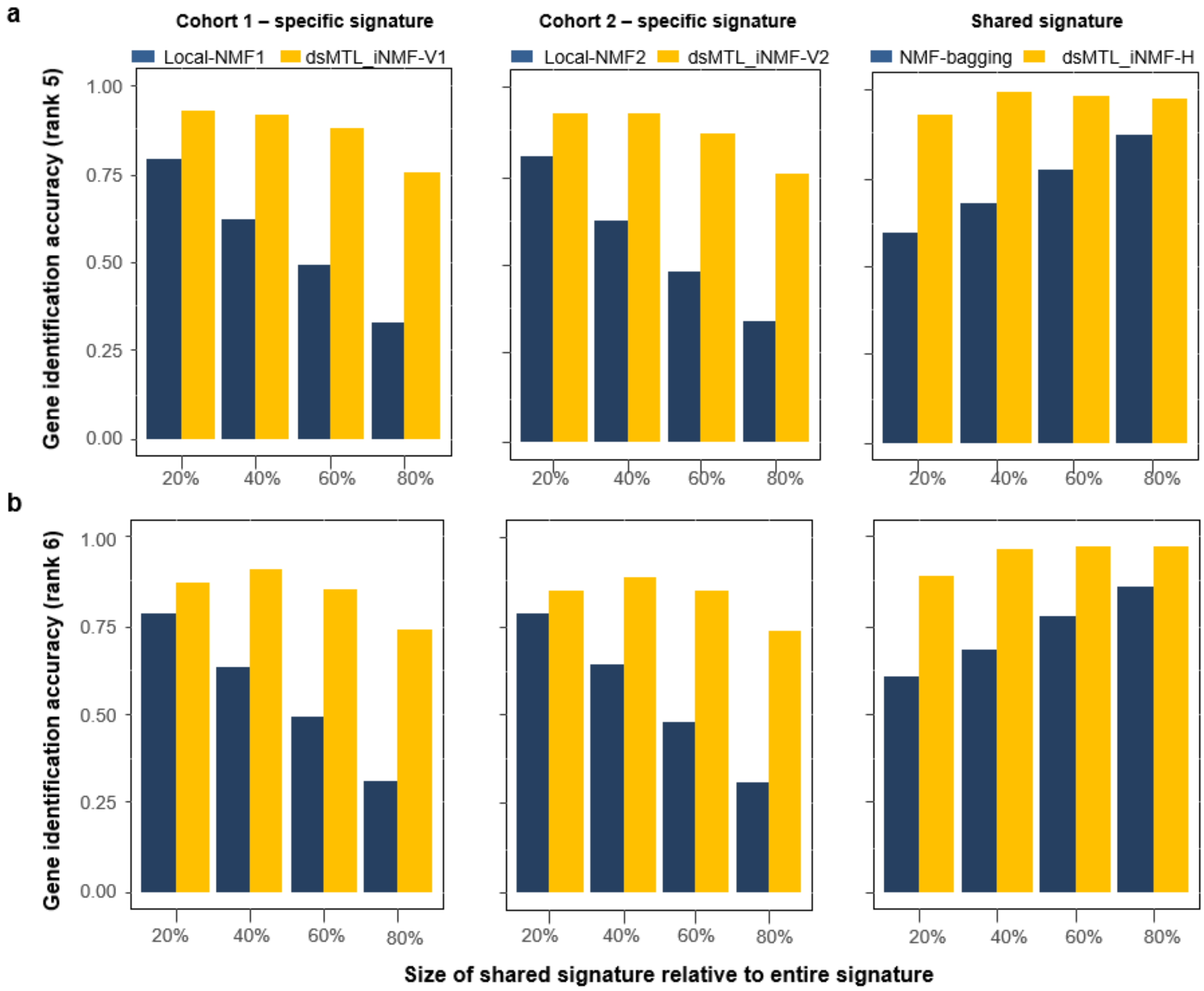


**Supplementary Figure 2. The gene identification accuracy for shared and specific signatures using simulated data using** rank=5 (**a**) and rank=6 (**b**) as model parameters.

| Index of test | Proportion of homogenous signatures | Number of samples | Number of features | Number of signatures | Proportion of patients |
|---|---|---|---|---|---|
| 1 | 20% | 200 | 1000 | 500 | 50% |
| 2 | 40% | | | | |
| 3 | 60% | | | | |
| 4 | 80% | | | | |

**Supplementary Table 1.** The simulation data of each server. These parameters were same to each of two servers. The only difference is the set of heterogeneous signatures.

| | | Server 1 | Server 2 | Server 3 | Client |
|---|---|---|---|---|---|
| Type | | Training | Training | Training | Testing |
| Location | | Mannheim | Mannheim | Heidelberg | Mannheim |
| Hardware | CPU | I7-4790 3.6GHz | I7-4790 3.6GHz | Intel Xeon 2.4 GHz | I7-4790 3.6GHz |
| | Memory | 4G | 4G | 4G | 16G |
| ID | | GSE35977 | GSE21138 | GSE53987 | HBCC |
| Number of Subjects | | 101 | 59 | 34 | 422 |
| Number of Genes | | 8013 | 8013 | 8013 | 8013 |

**Supplementary Table 2.** Client-server architecture for the real data analysis.

| | | dsLasso | Mean regularized dsMTL |
|---|---|---|---|
| Misclassification rate | | 0.34 | 0.29 |
| Time consumed | 5-fold CV | 7.5 mins | 7.3 mins |
| | Training | 1.7 mins | 2.9 mins |
| Number of network accesses for training | | 70 | 137 |
| Non-zero coefficients | | 38 | 173 |

**Supplementary Table 3.** Performance of dsML/MTL models on real data in real network.

| | | Server 1 | Server 2 |
|---|---|---|---|
| Type | | Training | Training |
| Location | | Heidelberg | Mannheim |
| Hardware | CPU | Intel Xeon 2.4 GHz | I7-4790 3.6GHz |
| | Memory | 4G | 4G |
| ID | | GSE164376 | GSE134497 |
| Number of Subjects | | 17 | 16 |
| Number of Genes | | 15215 | 15215 |

**Supplementary Table 4.** Details of server configurations used for real data analysis.
